# MTF1, a classic metal sensing transcription factor, promotes myogenesis in response to copper

**DOI:** 10.1101/534271

**Authors:** Cristina Tavera-Montañez, Sarah J. Hainer, Daniella Cangussu, Shellaina J.V. Gordon, Yao Xiao, Pablo Reyes-Gutierrez, Anthony N. Imbalzano, Juan G. Navea, Thomas G. Fazzio, Teresita Padilla-Benavides

## Abstract

MTF1 is a conserved metal-binding transcription factor in eukaryotes that binds to conserved DNA sequence motifs, termed metal response elements (MREs). MTF1 responds to metal excess and deprivation, protects cells from oxidative and hypoxic stresses, and is required for embryonic development in vertebrates. We used multiple strategies to identify an unappreciated role for MTF1 and copper (Cu) in cell differentiation. Upon initiation of myogenesis from primary myoblasts, MTF1 expression increased, as did nuclear localization. *Mtf1* knockdown impaired differentiation, while addition of non-toxic concentrations of Cu^+^ enhanced MTF1 expression and promoted myogenesis. Cu^+^ bound stoichiometrically to a C-terminus tetra-cysteine of MTF1. MTF1 bound to chromatin at the promoter regions of myogenic genes and binding was stimulated by copper. MTF1 formed a complex with MyoD at myogenic promoters, the master transcriptional regulator of the myogenic lineage. These studies establish novel mechanisms by which copper and MTF1 regulate gene expression in myoblast differentiation.

## Introduction

Copper (Cu) is an essential micronutrient required for human development and function. Cu plays a role in several key cellular functions, such as respiration, antioxidant defense, neurotransmitter biogenesis, disproportionation of O^−^, and metal ion homeostasis [1–3]. Dysregulation of cellular Cu levels is detrimental to human health, and it is associated with redox stress, disruption of iron-sulfur cluster proteins, lipid peroxidation, and DNA oxidation [4]. Consequently, cells must control Cu levels and prevent the accumulation of labile Cu in the cytosol. Cu homeostasis is maintained by a complex cellular network of transmembrane transport systems, soluble chaperones, chelating proteins, and transcription factors (TFs) [3, 5–8]. Cu depletion or overload leads to pathological conditions, such as Menkes’ disease and Wilson’s disease [8–15]. Menkes’ disease is characterized by severe Cu deficiency due to mutations in the Cu^+^-ATPase ATP7A that disrupt dietary Cu absorption. These inactivating mutations result in neurological abnormalities, blood vessel and connective tissue defects, and weak muscle tone (hypotonia) [16–20]. Wilson’s disease, which arises from mutations in the Cu^+^-ATPase ATP7B, results in Cu accumulation in the liver, brain, and eyes [19, 21, 22]. This Cu overload leads to a variety of hepatic and neurological defects, cardiomyopathies, and muscular abnormalities, such as a lack in coordination (ataxia) and repetitive movements (dystonia) [23, 24].

Copper is a fundamental co-factor for several enzymes, including cytochrome *c* oxidase (COX), and superoxide dismutases (SOD1 and SOD3) [1, 2]. Cu is also an important component of enzymes that contribute to proper tissue function [25–28]. Myogenesis encompasses several metabolic and morphological changes that are linked to Cu^+^-dependent cellular energy production and redox homeostasis [1, 2]. During myoblast differentiation, a metabolic shift occurs in which energy is produced via oxidative phosphorylation [29, 30]. This metabolic shift involves an increase in the production of mitochondria and associated cuproenzymes essential for energy production (*e.g*. COX) and redox homeostasis (*e.g*. SOD1) [1, 2, 30, 31]. Dysfunction or inhibition of mitochondrial protein synthesis impairs myogenesis [32–35].

We recently demonstrated that copper is required for the proliferation and differentiation of primary myoblasts derived from mouse satellite cells, which are adult stem cells responsible for skeletal muscle growth and repair from injury [36]. During myogenesis, the cellular levels of Cu increased, which is consistent with a high demand for Cu for proper function of mature tissue [36]. These changes in Cu levels are dependent on the dynamic expression of the Cu^+^-transporters and the post-transcriptional regulation of *Atp7a* [36]. However, the mechanisms by which Cu elicits a differentiation effect are unknown. Here, we hypothesized that Cu may have a fundamental role in the regulation of gene expression that drives the differentiation of the skeletal muscle. Activation of the myogenic program at the transcriptional level requires a series of signals, including growth factors, TFs, kinases, chromatin remodelers, histone modifiers, and metal ions [36–52]. Emerging evidence suggests that Cu and potential Cu^+^-binding TFs play significant roles in mammalian development [53–56]. Despite this, only three Cu^+^-binding factors are known to regulate gene expression in mammalian cells, and little is known about their roles in developmental processes [53, 54, 57–66].

Among these, MTF1 is a highly conserved zinc (Zn)-binding TF that recognizes and binds metal responsive elements (MREs) to promote the transcription of genes that maintain metal homeostasis [57, 59, 61, 67–70]. MREs are characterized by the -TGCRCNC-consensus sequence located near the promoters of genes related to redox and metal homeostasis [71–73]. MTF1 transcriptional activity is associated with the availability of Zn ions [74]; however, the molecular mechanisms by which metals activate MTF1 remain unclear. Current models for MTF1 activation include: (A) stimulation by free cytosolic Zn; (B) interaction with Zn released from metallothioneins (MTs); or (C) MTF1 phosphorylation/dephosphorylation [73, 75–78]. Under normal conditions, MTF1 is primarily located in the cytoplasm. When MTF1 is activated, it translocates from the cytoplasm to the nucleus, where it recognizes and interacts with MREs of genes that mediate homeostasis [61, 67, 79–84]. Chromatin immunoprecipitation (ChIP) analysis of *Drosophila* MTF1 has shown that different metal stimuli result in variations in the recognition of single nucleotides in genomic DNA sequences, showing that binding specificity can be altered by the presence of different metals [85]. *Drosophila* MTF1 has a Cu^+^ sensing function that is mediated in part by a carboxy-terminal tetra-nuclear Cu^+^ cluster [86]. A similar Cu^+^-binding centre has been identified in mammalian MTF1, suggesting that it may also respond to Cu [86]. Whether this response is associated with maintenance of metal homeostasis, or if it is related to other cellular functions, remains to be elucidated.

We have found that MTF1 is induced and translocated to the nucleus upon initiation of myogenesis in primary myoblasts derived from mouse satellite cells. shRNA and CRISPR/Cas9-mediated depletion of *Mtf1* causes lethality of differentiating myoblasts, indicating that MTF1 is essential for myogenesis. Nuclear levels of Cu increase in differentiating primary myoblasts and significantly decrease upon partial deletion of *Mtf1. In vitro* characterization of the murine MTF1 carboxy-terminal binding domain determined it bound stoichiometrically to Cu^+^. ChIP-seq and qPCR analyses revealed novel MTF1 target genes that are associated with myogenesis in addition to classic metal homeostasis genes. MTF1 interaction with myogenic genes is enhanced by supplementation of non-toxic concentrations of Cu to the myoblast differentiation media. Finally, our data indicate that one potential mechanism by which MTF1 participates in the transcriptional regulation of myogenic genes is through an interaction with MyoD. Taken together, our results shed light on the under-appreciated role of Cu and Cu-binding TFs in the development of skeletal muscle.

## Results

### MTF1 is up-regulated during differentiation of primary myoblasts

MTF1 is a metal binding transcription factor that is primarily involved in the control of metal and redox homeostasis [57, 59, 61, 67, 69, 70, 82, 83, 85, 87–91]. There is also evidence to suggest that MTF1 is involved in developmental processes [58, 62, 64, 92]. We hypothesized that MTF1 may play an active role in the determination of the myogenic lineage. To test this, we analysed both the expression and localization of MTF1 in primary myoblasts derived from mouse satellite cells. Western blot analyses showed minimal expression of the MTF1 protein in proliferating primary myoblasts (**Fig. 1A, B**). However, MTF1 protein expression was upregulated when differentiation was induced, as shown by the expression of myogenic markers (**Fig. 1A**). Confocal microscopy imaging of MTF1 is consistent with western blot analyses (**Fig. 1B**). Proliferating primary myoblasts have low levels of MTF1 in a punctate cytosolic distribution. Upon induction of myogenic differentiation, MTF1 expression increased and was primarily localized to the nucleus. At 48 and 72 h after initiation of differentiation, the distribution of MTF1 was primarily nuclear, though there was an increase in the cytosolic puncta that is consistent with its role in metal sensing. The data indicate that differentiation induces MTF1 expression and nuclear localization.

**Figure 1.**
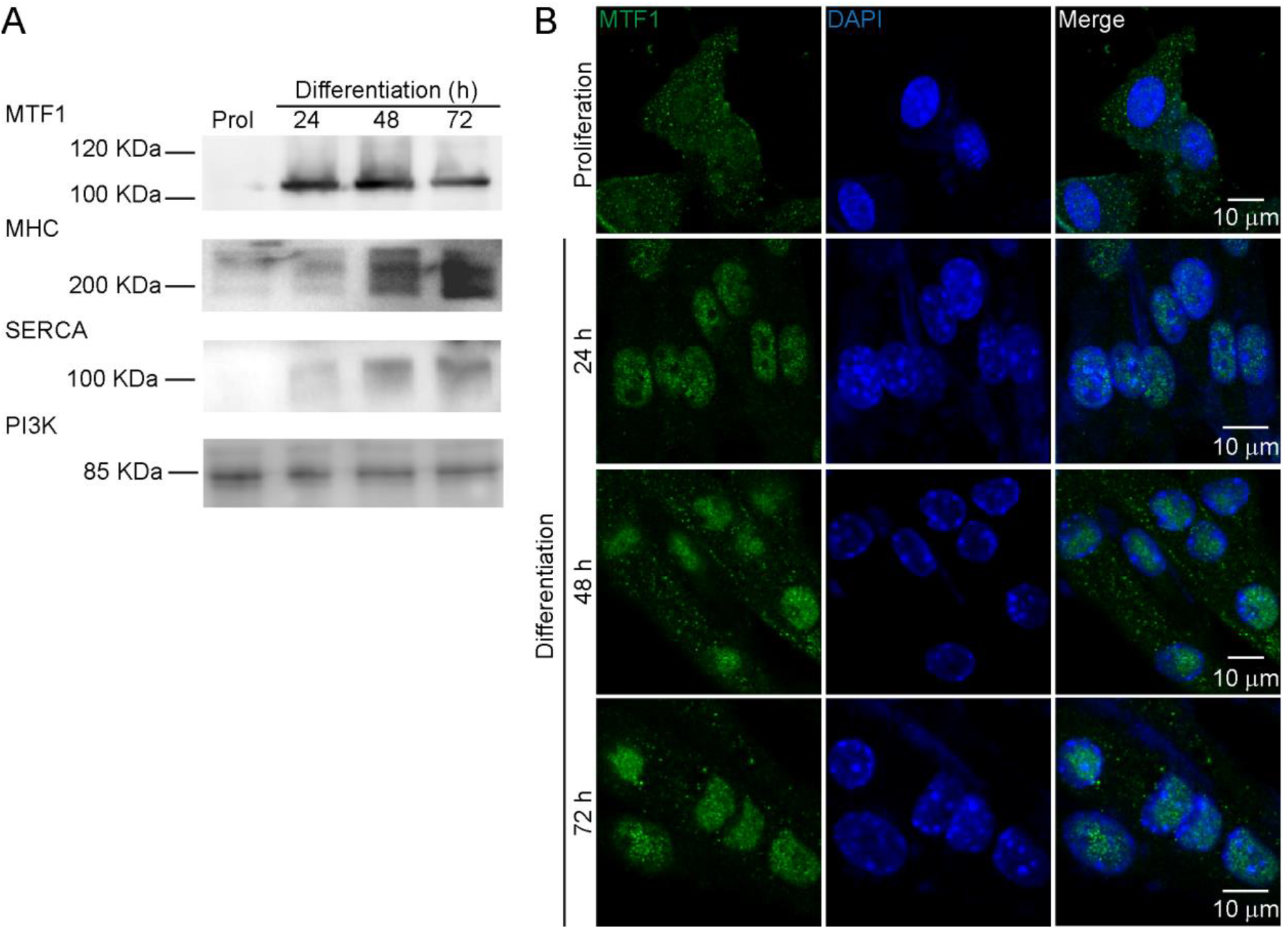
MTF1 is induced upon induction of differentiation of primary myoblasts. (A) Representative western blot of MTF1 expression; the myogenic differentiation markers examined were myosin heavy chain (MHC), and the sarco/endoplasmic reticulum Ca^2^+-ATPase (SERCA) from proliferating and differentiating primary myoblasts at 24, 48 and 72 h. PI3K was used as loading control. (B) Representative confocal microscopy images of proliferating and differentiating primary myoblasts at 24, 48 and 72 h for MTF1 (green), and DAPI (blue). Images depicted are representative of at least three independent biological experiments.

### *Mtf1* is required for myoblast differentiation

To determine the physiological role of MTF1, we used viral vectors encoding small hairpin RNAs (shRNA) to knock down *Mtf1*, and the CRISPR/Cas9 system to generate *Mtf1*-deficient primary myoblasts. Two lentiviral constructs that encode for shRNAs against *Mtf1* mRNA were used to knock down the endogenous protein in proliferating and differentiating primary myoblasts. Myoblasts transduced with a lentivirus-encoded non-specific scramble shRNA were used as negative controls. The virus-infected cells were selected with puromycin and the levels of *Mtf1* were examined by Western blot analysis (**Fig. 2**). Differentiating myoblasts transduced with *Mtf1* shRNA showed a decrease in the expression of MTF1 protein compared to wild type and *scr* shRNA controls (**Fig. 2A**). *Mtf1* knockdown cells had similar growth kinetics compared to wild type cells, suggesting that proliferating primary myoblasts can tolerate MTF1 knockdown (**Fig. 2B**). These results suggests that the primary role of MTF1 in proliferating myoblasts is maintenance or metal homeostasis as opposed to regulation of the cell cycle. To determine whether the partial loss of MTF1 impaired myogenesis, *Mtf1* shRNA-transduced primary myoblasts were induced to differentiate. Western blot analyses for the differentiation marker sarco/endoplasmic reticulum Ca^2+^-ATPase (SERCA) showed decreased levels in myoblasts partially depleted of MTF1 (**Fig. 2A**). Immunohistochemistry (IHC) analyses where differentiating myoblasts were stained with an anti-myogenin antibody confirmed that *Mtf1* knockdown cells fail to differentiate, as they had a decreased incidence of myogenin-positive nuclei compared to wild type and *scr* shRNA transduced myoblasts (**Fig. 2C**). Detachment of *Mtf1* knockdown cells was observed upon induction of differentiation (**Fig. 2C**). Western blot analyses showed an increased expression and activation of the apoptotic marker Caspase 3 in differentiating myoblasts partially depleted of *Mtf1* (**Fig. 2A**).

**Figure 2.**
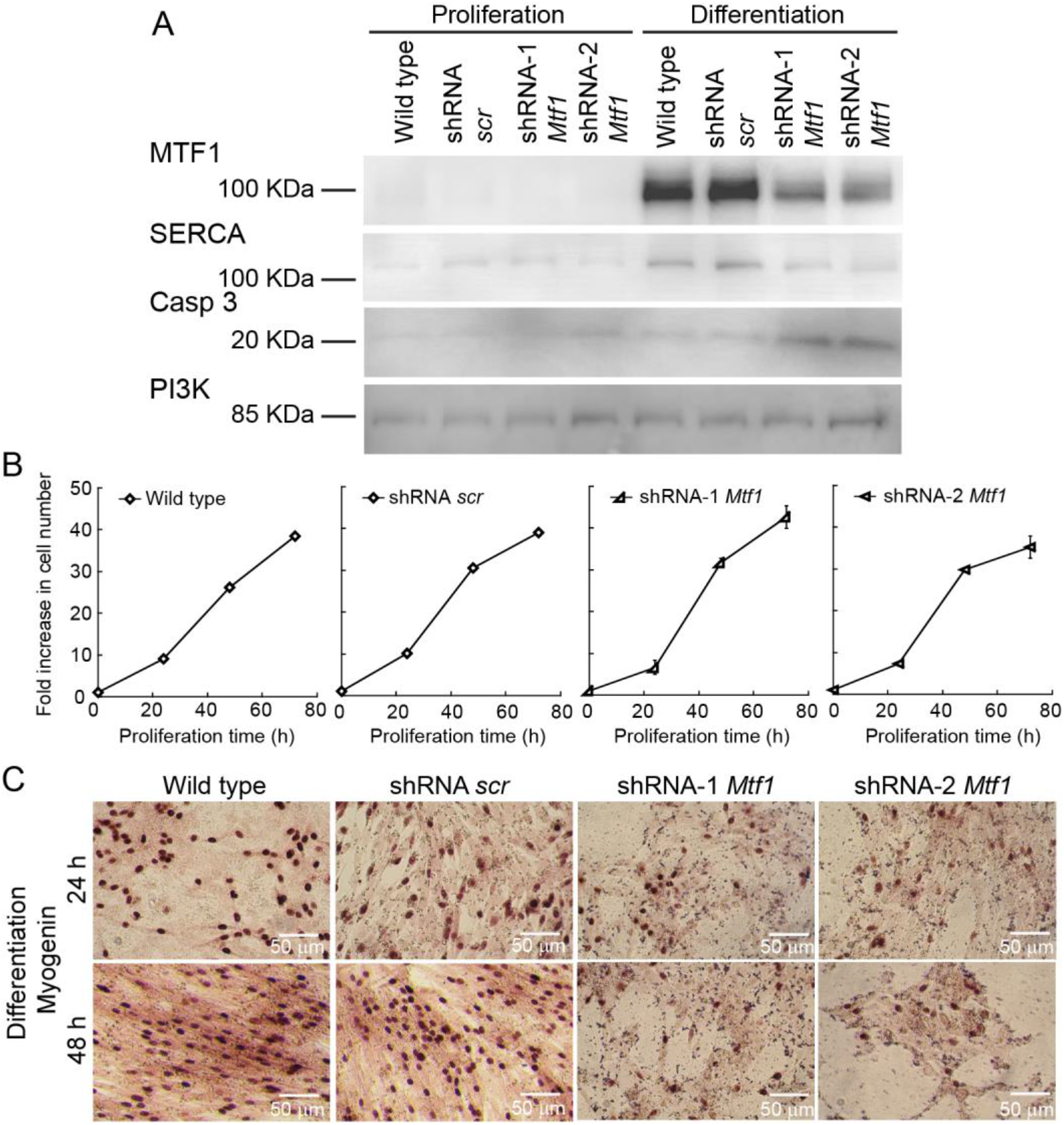
Partial depletion of MTF1 using shRNA impairs myogenesis and partially leads to death of differentiating myoblasts. (A) Representative Western blot of primary myoblasts and myoblasts transduced with either scramble shRNA (scr) or two different shRNAs against *Mtf1* (1 and 2) during proliferation and 24 h after inducing differentiation. SERCA levels were monitored as a differentiation marker, cleaved Caspase 3 as marker of cell death, and PI3K as a loading control. (B) Proliferation curves comparing wild type, scramble control (scr), and MTF1 (shRNA 1 and 2) partially depleted primary myoblasts. No significant differences were found between the four strains. Data represent the average of three independent experiments ± SE. (C) Representative light micrographs of differentiating myoblasts immunostained for myogenin at 24 and 48 h.

The biological relevance of MTF1 during myogenesis was confirmed by targeting the *Mtf1* locus with CRISPR/Cas9. Western blot analyses of differentiating primary myoblasts at 24 h showed over a 90% reduction in MTF1 protein and gene levels (**Fig. S1**). Consistent with our observations using shRNA (**Fig. 2**), MTF1 loss correlated with a failure to differentiate as shown by the decrease in protein and gene expression of myogenic markers (**Fig. S1A-B**). *Mtf1*-deficient cells proliferated normally and showed no visible phenotype during initial passages (**Fig. S1C**), however extended culture (5 passages) of these cells resulted in increased apoptosis relative to control cells (data not shown), suggesting that the MTF1-deficient cells are sensitive to extended passage in tissue culture. IHC analysis showed that *Mtf1*-deficient cells detached from the plates at 24 h after induction of myogenesis (**Fig. S1D**), which correlates with increased cleaved Caspase-3 (Casp3), as compared to wild type and empty vector sgRNA myoblasts (**Fig. S1A**). Overall, these results show that MTF1 plays an essential functional role in the regulation of the expression of myogenic genes and contributes to cell survival upon initiation of myogenic differentiation.

### MTF1 expression is enhanced by Cu ions

In order to induce myogenesis in cultured myoblasts, growth factors are depleted by serum starvation and insulin is added [93, 94]. Depletion of insulin from the differentiation medium partially prevents myogenic differentiation, a phenotype that we have recently shown can be rescued by the addition of non-toxic (30 μM) concentrations of CuSO_4_ [36]. Moreover, depletion of Cu from the culture medium inhibits differentiation, which suggests that Cu plays a role in differentiation [36]. However, the molecular mechanisms by which Cu affects differentiation are largely unknown. To probe for links between Cu ions and MTF1, we cultured primary myoblasts under different concentrations of Cu and determined the expression levels of MTF1 through Western blot and qPCR. **Figure 3** shows that MTF1 expression was significantly increased in cells grown in medium depleted of insulin and supplemented with 30 μM CuSO_4_ compared to those grown in basal differentiation medium containing insulin. By contrast, Cu chelation with tetraethylenepentamine (TEPA) resulted in a significant decrease in MTF1 expression. Addition of CuSO_4_ at a concentration equal to that of TEPA restored the expression levels of MTF1 to those observed in cells differentiated under normal insulin conditions. These data indicate a role for Cu ions in MTF1 induction during myogenesis. Importantly, we detected an increase in the expression of the differentiation marker SERCA when the cells were treated with Cu, which was abolished when the cells were depleted of Cu by addition of TEPA (**Fig. 3A**). These data are consistent with our previous studies where the expression of myogenin and other differentiation markers was enhanced by copper supplementation [36].

**Figure 3.**
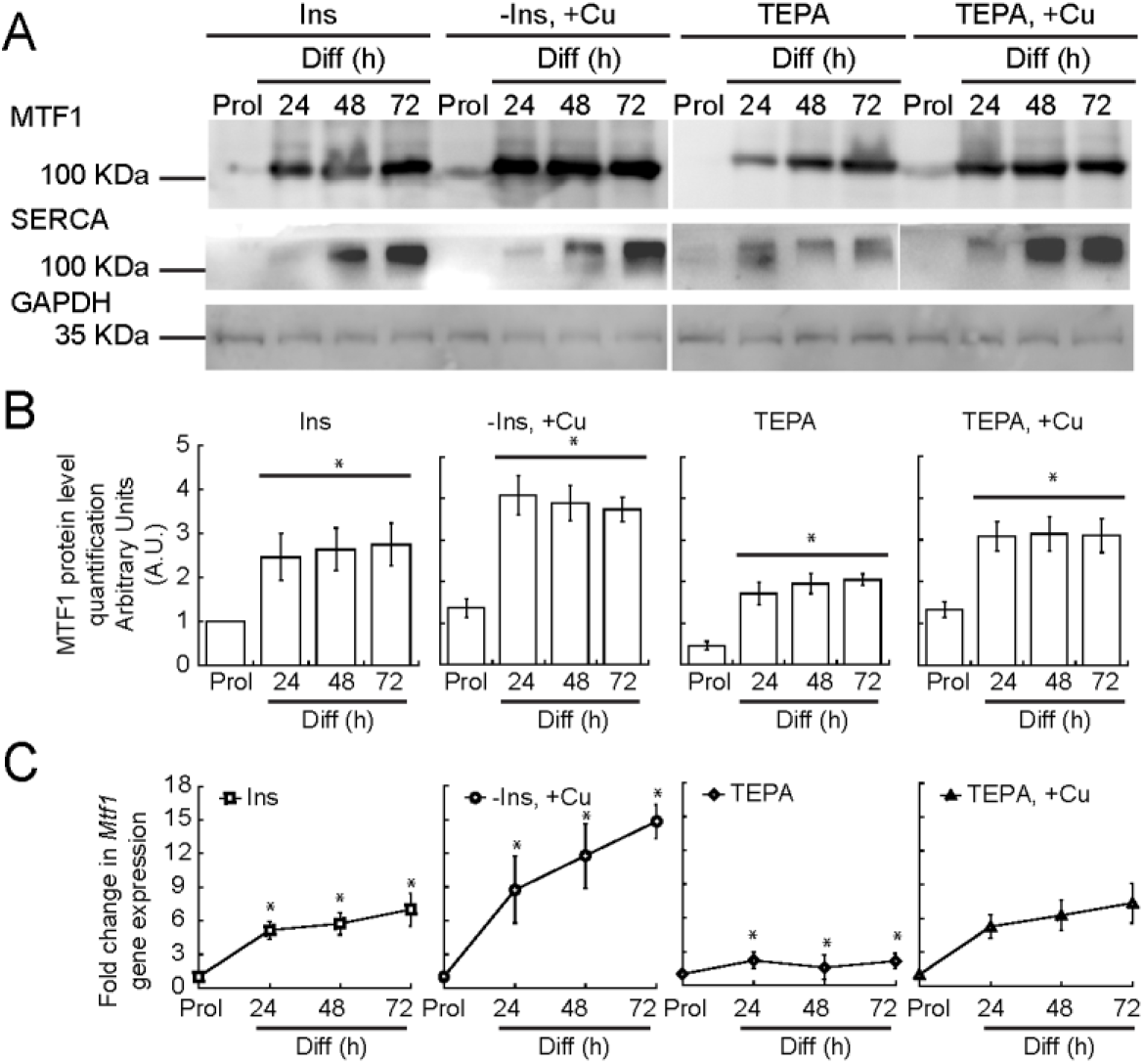
Copper enhances the expression of MTF1 in differentiating myoblasts. (A) Representative Western blots of MTF1 and SERCA in proliferating and differentiating myoblasts differentiated in the presence or absence of insulin, Cu and TEPA as indicated. GAPDH was used as loading control. (B) Densitometric quantification of MTF1 bands in proliferating and differentiating (24, 48 and 72 h) primary myoblasts. (C) Steady-state mRNA levels of *Mtf1* determined by qRT-PCR from proliferating and differentiating primary myoblasts cultured in the same conditions described in (A). The data represent the average of three independent biological experiments ± SE; *P < 0.01.

### MTF1 binds to the promoters of myogenic genes in differentiating myoblasts

Our data thus far indicate that MTF1 is required for myogenesis in addition to its role in maintaining metal homeostasis. We hypothesized that copper enhances the transcriptional activity of MTF1, and that MTF1 globally regulates the expression of genes required for skeletal muscle differentiation. To test this, we performed ChIP-seq to identify MTF1 binding sites on chromatin in primary myoblasts differentiated for 24 h under normal insulin conditions. Genome wide analyses showed that in the presence of insulin, MTF1 localized largely to promoters around transcriptional start sites (TSS; **Fig. 4A**). MTF1 binding to TSS was enhanced by the addition of CuSO_4_ to the culture media. Cells depleted of insulin from the culture media had decreased MTF1 binding that was similar to the input controls (**Fig. 4A**).

**Figure 4.**
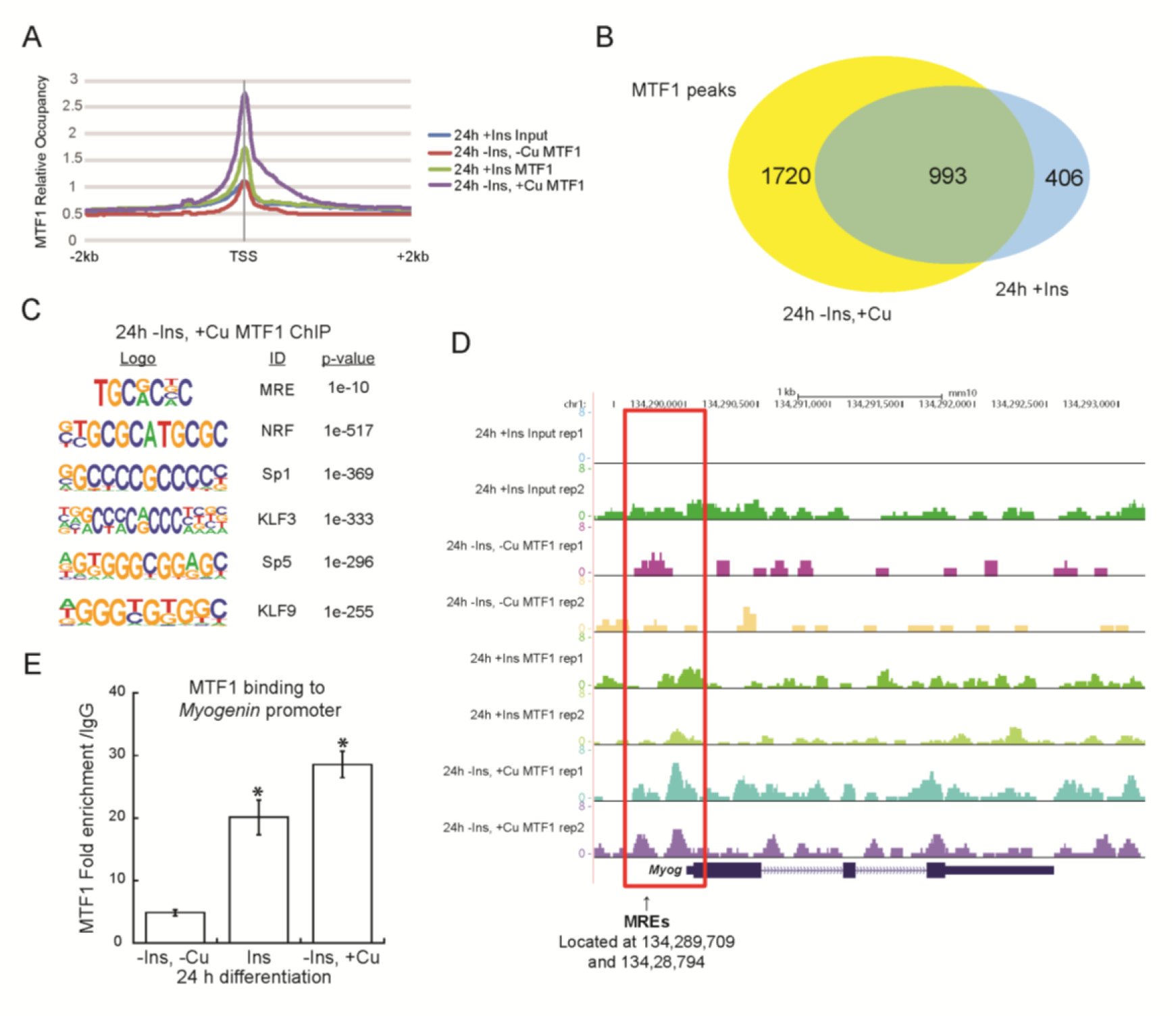
MTF1 binding in differentiating primary myoblasts. (A) Aggregation plots of MTF1 ChIP-Seq data showing occupancy over annotated TSS. (B) Overlap of ChIP-Seq peaks of MTF1 across the genome observed in differentiating cells in the presence of insulin or Cu. (C) Novel consensus DNA-binding motifs identified within MTF1 peaks. Shown are MRE and the top five most significant motifs enriched, including the DNA logo, its corresponding TF, and its p-value. (D) Genome browser tracks of replicate ChIP-Seq experiments examining MTF1 binding to the Myogenin promoter in differentiating myoblasts (24 h) under different culture conditions. (E) ChIP-qPCR validation for MTF1 binding to the Myogenin promoter. Data represent the average of three independent biological replicates ± SE; *P < 0.01.

Next, we called peaks for MTF1 in differentiating myoblasts. We found 1399 MTF1 peaks for myoblasts differentiated under normal insulin conditions, 2713 for cells differentiated in the presence of Cu, and only 553 peaks for the cells depleted of insulin. Strikingly, 993 peaks were shared between the cells differentiated with insulin and the cells differentiated with Cu (**Fig. 4B, Table S1**). Gene ontology (GO) analyses showed that MTF1 binds to diverse categories of genes associated with muscle development, function and as expected, ion homeostasis (**Fig. S2; Table S2**).

MTF1 is known to preferentially bind the MRE consensus sequence, -TGCRCNC-[71–73]. As expected, ChIP-seq analyses showed MTF1 enriched binding to this motif (**Fig. 4C, Table S1**). In the presence of insulin or CuSO_4_, 32% and 38% of the MTF1 peaks contained a consensus MRE, respectively. This result suggests direct binding by MTF1 to MREs as one mechanism of MTF1-mediated gene activation, but also suggests that MTF1 is interacting with chromatin indirectly through other TF binding sites. We performed a *de novo* motif search on the Cu MTF1 peaks to identify additional binding motifs for MTF1. The regions under MTF1 peaks had high GC contents, and the top five motifs are shown (**Fig. 4C, Table S1**). The most significant of these motifs matches to the binding motif for the TF nuclear respiratory factor (NRF). NRF-1 and −2 play a role in the expression of nuclear and mitochondrial genes involved in oxidative phosphorylation, electron transport complexes I-V, and mtDNA transcription and replication [95, 96], all processes important in skeletal muscle function. The second most significant motif matches the binding site of Sp1, a TF that has been shown to regulate muscle gene expression in concert with MyoD [97]. The next most significant motifs correspond to KLF3, SP5, and KLF9, each of which has been linked to skeletal muscle differentiation or function [98–101]. These data suggest that MTF1 may bind chromatin in conjunction with multiple diverse regulators to regulate gene expression during muscle differentiation.

We found the conserved MRE sequence for MTF1 binding at some myogenic genes, such as *Myogenin*, which encodes the classic transcriptional activator that is expressed upon initiation of myogenesis and is essential for the transcription of muscle-specific genes [41, 48, 102, 103]. Bioinformatic analyses revealed that two potential MREs, 5’-TGCACAG-3’ and 5’-TGCACCC-3’, are located at 300 and 400 base pairs upstream, respectively, of the *Myogenin* TSS. Therefore, we hypothesized that MTF1 may bind near the promoter of *Myogenin*. To test this, we assessed our ChIP-seq data over the *Myogenin* promoter and found that MTF1 binding was enriched by Cu supplementation (**Fig. 4D**). The binding of MTF1 to *Myogenin* was validated by ChIP-qPCR, which showed a significant increase in binding when the myoblasts are differentiated in media supplemented with 30 μM CuSO_4_ (**Fig. 4E**).

Binding of MTF1 was evaluated at additional myogenic genes by ChIP-seq and ChIP-qPCR. There is enhanced binding of MTF1 to the promoter of <u>A</u> <u>D</u>isintegrin <u>A</u>nd <u>M</u>etalloproteinase 9 (*Adam9*), a membrane anchored cell surface adhesion protein that mediates cell-cell and cell-matrix interactions and muscle development (**Fig. S3A, B**) [104]. MTF1 was also found at the promoters of additional myogenic genes, such as *MyoD, Integrin 7a, Skeletal actin, Myf5* and *Cadherin 15* (**Table S2**). To validate our ChIP-seq analyses, we analyzed MTF1 binding to its classic target promoter, *Metallothionein 1* (*Mt1*). ChIP-qPCR data showed that MTF1 binding to the *Mt1* promoter is enhanced upon addition of 30 μM CuSO_4_ to the culture media of primary myoblasts, as expected (**Fig. S3C, D**). As a negative control, no binding to the *Pax7* promoter was observed (**Fig. S4**).

### MTF1 interacts with MyoD and binds a subset of MyoD-bound loci

Interestingly, the classic DNA binding motif (E-box) of MyoD was included among the TF binding sites found within MTF1 peaks, suggesting a potential novel interaction at promoter regions for both transcription factors (**Table S1, line 33**). MyoD and MyoD-related factors initiate the regulation of skeletal muscle gene expression through direct binding of the promoters of myogenic genes during differentiation [97, 105]. We hypothesized that MTF1 may interact with MyoD, forming a complex that binds the promoters of myogenic genes that MyoD regulates. We first investigated whether MTF1 and MyoD physically interact in primary myoblasts differentiated for 24 h. Immunoprecipitation (IP) assays using an anti-MTF1 antibody revealed that MTF1 co-precipitated with MyoD upon initiation of myogenesis in both the presence and absence of Cu (**Fig. 5A**). To further characterize the functional relationship between MTF1 and MyoD, we compared the ChIP-seq datasets for MyoD in differentiated primary myoblasts (10 and 48 h) from Soleimani *et al*., [106] with our MTF1 ChIP-seq data. We found 714 peaks shared between MyoD and MTF1 when the myoblasts were differentiated in the presence of Cu, which represents over 25% of the total number of MTF1 peaks (**Fig. 4B**). GO term analyses of these peaks showed MyoD and MTF1 bind to myogenic genes, but also metal ion transport and homeostasis genes (**Fig. S2, Table S3**). *De novo* motif identification of overlapping MyoD and MTF1 peaks gives a similar outcome as the analysis of motifs under MTF1 peaks (**Fig. 5C, 4C**). Analysis of individual genes for MTF1 and MyoD binding showed increased peaks at the same promoter region of the *Myogenin* gene (**Fig. 5D**). This co-binding was confirmed by sequential re-ChIP analyses of MyoD and MTF1, which indicated that both TFs co-occupy the *Myogenin* promoter in myoblasts (**Fig. 5E**). We did not detect co-binding of the MTF1-MyoD complex to the *Pax7* promoter region (**Fig. 5F**), which further supports our conclusion that MTF1 regulates differentiation-specific gene expression. Together, these data suggest that MyoD and MTF1 form a stable complex on chromatin in differentiating primary myoblasts to regulate the transcription of a common set of myogenic target genes.

**Figure 5.**
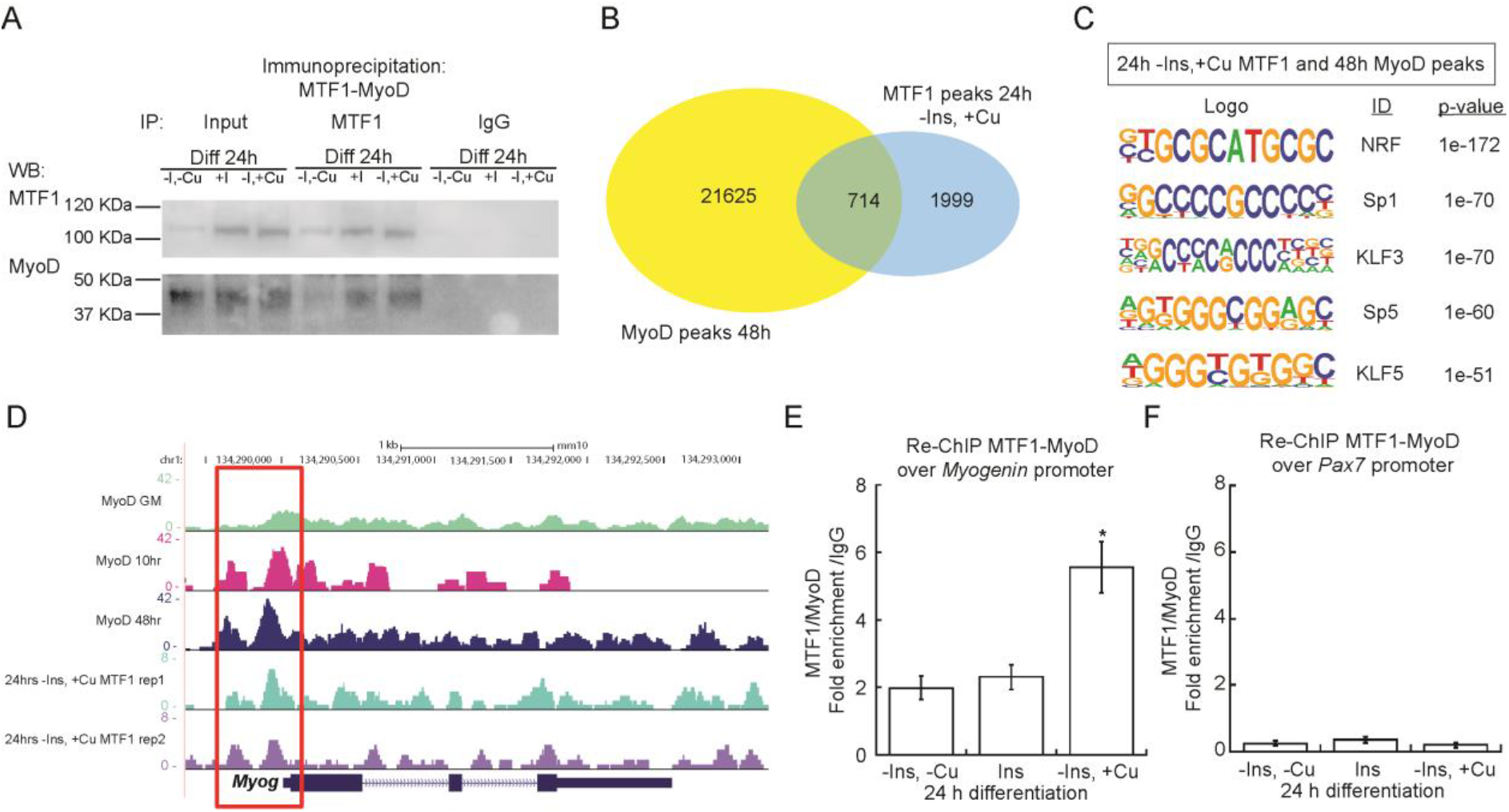
MTF1 interacts with MyoD at the promoter regions of myogenic genes. Representative Western blot of MTF1-MyoD immunoprecipitation (IP). Pulldown with IgG was used as a negative control. (B) Overlap of MTF1 ChIP-seq peaks from primary myoblasts differentiated with Cu MTF1 and MyoD peaks extracted from datasets published by Solemaini et al., 2012 (GSE24852). (C) Consensus binding motifs identified for sequences bound by MTF1 and MyoD. Shown are the top five most significant motifs enriched, including the DNA logo, its corresponding TF, and its p-value. (D) Genome browser tracks of ChIP-seq data comparing MTF1 occupancy at the myogenin locus in cells 24 h after differentiation with Cu and MyoD occupancy at the myogenin locus in proliferating (GM) cells or cells differentiated for 10 or 48 hours. MyoD ChIP-seq data was downloaded and analyzed from GSE24852 (Soleimani et al., 2012) [106]. (E-F) Reciprocal chromatin immunoprecipitation (Re-ChIP)-qPCR for MTF1-MyoD co-binding to the Myogenin promoter (E) and to the Pax7 promoter (F) as a negative control. Data represent the average of three independent biological replicates ± SE; *P < 0.01.

### MTF1 binds Cu, which may play a role in its nuclear translocation and enhanced binding to myogenic promoters

We recently reported that differentiating myoblasts accumulate Cu, which is consistent with the inherent requirement for this metal during myogenesis [36]. Consistent with this hypothesis, atomic absorbance spectroscopy (AAS) analyses showed that the increase in Cu levels observed in differentiating myoblasts is prevented upon Cu chelation (**Fig. S5A**). Subcellular fractionation of proliferating and differentiating myoblasts showed that Cu is mobilized to the nucleus upon induction of myogenesis (**Fig. S5B**). Cells grown in the presence of Cu had higher levels of Cu in the nucleus than did control cells during proliferation. Lower levels of nuclear Cu were detected in myoblasts differentiated in the presence of TEPA (**Fig. S5B**). Cytosolic concentrations of Cu were higher than in the nucleus under all conditions tested (**Fig. S5C**).

We hypothesized that the potential of MTF1 to bind copper may contribute to the nuclear translocation and enhanced activation of this TF. Interestingly, *Drosophila* MTF1 contains a carboxy-terminal cysteine-rich Cu^+^-binding domain, distinct from its Zn finger domains that bind Zn ions, that is proposed to sense excess intracellular Cu and participate in the cellular heavy metal response [75, 86]. Mammalian MTF1 has a similar putative Cu^+^-binding domain at its C-terminal domain. To test for a similar function, we cloned, expressed, and purified wild type murine recombinant MTF1 (**Fig. 6A**). We first examined the Cu^+^-binding properties of the purified protein by incubating it with excess Cu^+^ in the presence of ascorbate as reducing agent. Metal determinations by AAS revealed that Cu^+^ interacts with MTF1 at a stoichiometry of 1.17 ± 0.06 Cu atoms per protein (**Fig. 6B**). To test whether Cu binds at the C-terminal cysteine-rich domain, we mutated the four key cysteine residues to alanines (metal binding site (MBS), **Fig. 6A**). These mutations strongly impaired the binding of Cu^+^ to MTF1 (**Fig. 6B**), implicating these amino acids in Cu^+^ binding.

These data raise the possibility that MTF1 contributes to the translocation of Cu ions to the nucleus. To test this hypothesis, we took advantage of primary myoblasts partially depleted of *Mtf1* by shRNA (**Fig. 2**). We evaluated whether MTF1 knockdown would impair the capability of the cells to translocate Cu into the nucleus. Overall, *Mtf1* knockdown cells exhibited a significant decrease in the total levels of Cu upon induction of differentiation, whereas only a non-significant, but consistent small decrease in the levels of Cu in proliferating myoblasts was detected (**Fig. 6C**). Subcellular fractionation showed that *Mtf1* knockdown differentially affected the nuclear Cu content. For instance, proliferating *Mtf1* knockdown myoblasts contain only ~10% of nuclear Cu compared to wild type and *scr* controls. Nuclear fractions of differentiating *Mtf1* knockdown cells contained ~50% of Cu compared to the control cells (**Fig. 6D**). Minor changes in the cytosolic levels of proliferating and differentiating *Mtf1* knockdown myoblasts were detected (**Fig. 6E**). These data are consistent with a role of MTF1 mobilizing Cu into the nucleus during myogenesis, but does not exclude the possibility that other proteins are also part of this process.

**Figure 6.**
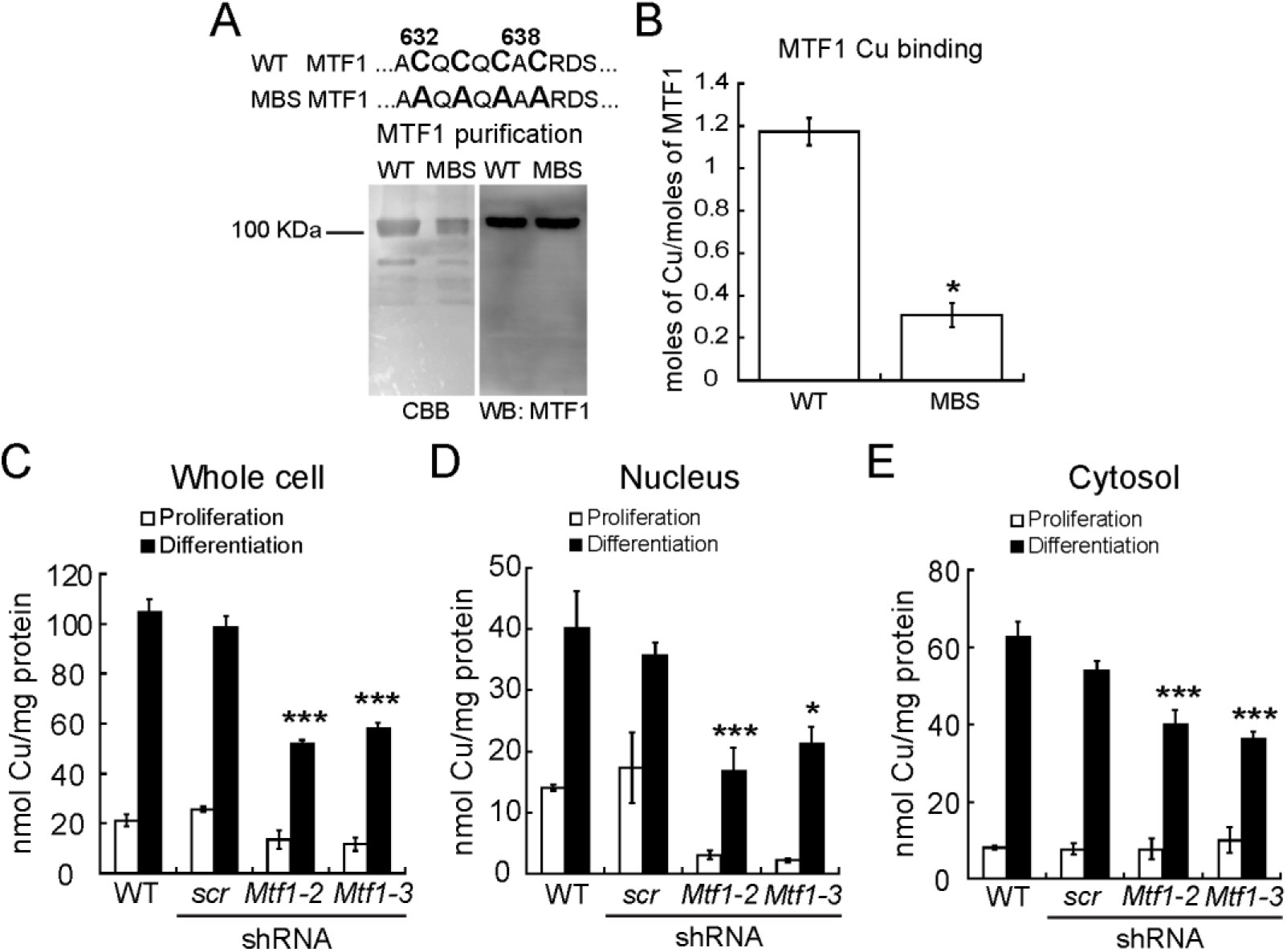
MTF1 binds Cu, which correlates with nuclear translocation of the metal. (A) Upper panel depicts the sequence containing the tetra-cysteine cluster at the carboxy-terminal of the murine MTF1 that is required for transcriptional response to Zn and Cd (amino acids 632, 634, 636 and 638) [86]. These residues were mutated to Alanine to asses Cu-binding capabilities of this putative <u>m</u>etal-<u>b</u>inding <u>s</u>ite (MBS). Lower panel shows a representative Coomassie Brilliant Blue-stained SDS/PAGE and a Western blot immunostained with anti-MTF1. (B) Cu binding stoichiometry of purified wild type and MTF mutated at the Cu-binding site (MBS) determined by AAS. (C) Whole cell Cu content of proliferating and differentiating primary myoblasts determined by AAS. Nuclear (D) and cytosolic (E) Cu content of proliferating and differentiating primary myoblasts cultured under different Cu conditions. Metal determination was performed by AAS. Data represent the mean Cu concentration of three independent biological replicates ± SE.

## Discussion

There is a significant gap in our understanding of the roles that Cu plays in transcriptional regulation during mammalian development. Previous studies from our laboratory have shown that Cu promotes the proliferation and differentiation of primary myoblasts derived from mouse satellite cells [36]. The pathways and mechanisms by which this transition metal induces this myogenic effect are largely unknown. In this work, we characterized the roles of copper and the Cu-binding transcription factor, MTF1, in myogenesis. Our data shows that MTF1 expression is essential for myogenesis, and that Cu enhances the expression of MTF1. Moreover, we have found that cellular Cu content influences the binding of MTF1 to target promoters. Finally, our studies revealed multiple mechanisms of MTF1 interaction at target genes, including direct binding to MREs and presumed indirect interactions through other transcription factors, including MyoD.

MTF1 is activated by different mechanisms to control metal and redox homeostasis, which include stimulation by cytosolic Zn and/or Zn released from MTs, or regulation by phosphorylation events [73, 75–78]. On the other hand, the mechanisms by which MTF1 stimulates transcription of metal responsive genes (MTs and metal transporters) in response to heavy metals and oxidative stress is well-established. A characteristic of the promoters and enhancers of most of MTF1 target genes is the presence of MREs in the upstream regulatory sequences or just downstream of the TSS of metal-responsive genes that mediate MTF1 binding and regulation of gene expression [64, 71, 78, 83, 107].

Activation of MTF1 by copper has been investigated in several vertebrate models. Studies have addressed the expression of MT1 as an indirect measure of MTF1 activity upon stress induced by copper and other metals. For instance, *in vivo* studies showing Cu-dependent changes in the transcription of *Mt1* in mouse liver showed that only high doses (over 5 mg/kg) of Cu administered to the animals induced the expression of this gene, while little effect of Cu was detected in the kidney [108]. Studies in HeLa S3 cells showed that the transcript levels of the metal protective *MtIIA* gene increased upon treatment with 300 μM CuSO_4_. However, *in vitro* studies using whole cell extracts obtained from HeLa S3 cells exposed to 300 μM CuSO_4_ failed to induce the MRE-binding activity, attributed to MTF1, although this concentration of Cu was able to induce the expression of *MtIIA* [109]. In embryonic stem cells, induction of MT1 and MT2 was only achieved when the cells were treated with 500 μM CuSO_4_ [67]. It is noteworthy that studies from our laboratory showed that in differentiating, serum deprived, primary myoblasts derived from mouse satellite cells, concentrations over 100 μM are toxic to the cells [36]. The data presented here showed that supplementation of differentiating myoblast with 30 μM CuSO_4_, is sufficient to induce MTF1 expression and activation, not only to drive the expression of metal-protective genes, but also to promote the expression of myogenic genes. Overall, the data suggests that the cellular metal response and activation of MTF1 is dependent on the cell lineage. Studies from different laboratories suggest that copper treatment is a poor activator of MTF1 in the context of its classic metal protective role, as shown by the expression of *Mt1*. However, the studies shown here suggest that low concentrations of Cu contribute to the activation of MTF1 in a novel role as a modulator of the expression of genes associated with myogenesis. Studies should be now directed to investigate the roles of MTF1 in the development of other lineages.

Additional roles for MTF1 have been proposed during embryonic development [58]. MTF1 knockout leads to embryonic lethality at embryonic day 14 due to liver degeneration [58, 62]. The MTF1 target genes MT1 and MT2 are constitutively and highly expressed in fetal liver [110–112], suggesting that these proteins are fundamental for liver development. However, deletions of both metallothionein genes had no effect in development under normal conditions, but mice were sensitive to Cd stress [113, 114]. It is noteworthy that the MTF1 knockout murine model had no evidence of muscular phenotypes in developing embryos at E14 [58]. These findings suggest that 1) MTF1 contributes but is not required for muscle development; 2) that MTF1 contributes to early muscle development but is only required for developmental stages at or after E14, or 3) that there is an as yet unidentified redundancy for the roles of MTF1 in myogenesis. We also note an intriguing link between MTF1, Parkinson’s disease, and muscle function in *Drosophila*. Parkin is an E3 ubiquitin ligase mutated in some hereditary forms of Parkinson’s disease. The *Drosophila* homozygous mutant of the human ortholog *parkin* exhibits severe movement impairment, inability to fly, sterility, and short life span [92]. Overexpression of MTF1 in the *parkin* mutant flies partially rescues these phenotypes, likely due to an effect at the muscular and mitochondrial levels, supporting a role for MTF1 in muscle development [92]. In addition, myogenic regulatory factors such as MyoD, Myogenin, and the Myocyte Enhancer Factor 2 (MEF2) have been shown to regulate the expression of MTF1 in differentiating myoblasts, however no characterization has been done [115]. Therefore, the specific roles for MTF1 in development and in lineage determination remain to be elucidated. Our work suggests three potential mechanisms for MTF1 binding to myogenic genes: 1) Direct recognition and binding to MREs; 2) indirect binding through additional transcription factor binding sites; and 3) indirect binding through MyoD binding sites.

Studies in *Drosophila* MTF1 described a novel carboxy-terminal tetra-nuclear Cu center, independent of the Zn binding fingers [86]. Functional studies of the mammalian MTF1 showed a carboxy-terminal 13 amino acid domain that includes four conserved cysteines (CQCQCAC) that are necessary for MTF1 Zn and Cd sensing and transcriptional activation *in vivo* under moderate metal stress [116, 117]. Interestingly, these *Drosophila* and mammalian cysteine domains lack significant sequence homology and have been proposed to be the result of convergent evolution events of a Cu sensing element [86]. This cysteine cluster also mediates the homo-dimerization of MTF1, which is proposed to constitute a platform for the recruitment of additional transcriptional cofactors [118]. In this regard, Cu has been proposed to play a relevant role in stabilizing the dimer by constituting intermolecular disulphide bonds through oxidation of cysteines to further synergize with zinc to enhance transcription [118]. However, the mechanistic role of this domain in regulation of metal homeostasis and development genes is not yet clear. Importantly, our data corroborates that Cu is internalized during myoblast differentiation and a fraction of the internalized Cu is re-localized to the nuclei. Considering the significant increase in MTF1 expression upon addition of Cu and the Cu-binding capabilities of the carboxy-terminal cysteine cluster of MTF1, it is plausible that MTF1 is partially responsible for the nuclear translocation of copper observed in differentiating myoblasts. Studies are under way to further characterize the mechanistic roles of this novel Cu-binding domain in the regulation of gene expression associated with the myogenic lineage, metal and redox homeostasis, as well as the potential interactions between MTF1 and additional metal-dependent protein-protein interactions at MREs in promoter regions.

Current investigations in the field have been directed towards understanding the deleterious effects of Cu on the nervous system, liver, and intestine. However, little attention has been given to other organs and tissues, such as muscle, adipose, and bone. Strikingly, most of the systemic phenotypes observed in Menkes’ and Wilson’s disease patients have been attributed to the neurological damage that Cu exerts as a result from deficient systemic transport, rather than a direct effect on the different tissues and organs. However, a complex developmental process such as myogenesis encompasses metabolic and morphological changes linked to Cu-dependent energy production and redox homeostasis [1, 2]. Myoblast differentiation requires a metabolic shift in which energy is produced via oxidative phosphorylation, a process highly dependent on increased production of mitochondria, Cu availability and expression of cuproenzymes essential for energy production (e.g. COX) and redox homeostasis (e.g. SOD1) [1, 2, 30, 31]. Our results shed light onto the importance of the function of Cu and MTF1 in the regulation of gene expression during developmental processes, such as skeletal muscle differentiation. A better understanding of how tissue Cu status affects growth and development at other cellular levels, will be beneficial in the study of muscular phenotypes that present in diseases of Cu misbalance, such as Menkes’ and Wilson’s diseases.

## Material and methods

### Primary Cell Culture

Mouse satellite cells were isolated from leg muscle of 3-6 week old wild type C57Bl/6 mice. The muscle was extracted and cut into small pieces, washed with Hank’s Balanced Salt Solution (HBSS; ThermoFisher Scientific) and incubated with 0.1% Pronase for 1 h at 37 °C. The cells were then filtered using a 100 μm cell sieve and resuspended in 3 ml of growth media (1:1 v/v DMEM:F-12, 20% FBS, and 25 ng/ml of basic FGF). Cells were filtered again using a 40 μm cell sieve and centrifuged at 1,000 x *g* for 1 min at room temperature. The cells were placed at the top of a Percoll step-gradient (35 and 70%) and centrifuged 20 min at 1,850 x *g* at room temperature. The myoblasts were contained in the lower interface of the 70% Percoll fraction and were washed with HBSS, centrifuged 5 min at 1,000 x g, and resuspended in growth media for plating. Myoblasts were grown and differentiated on plates coated with 0.02% collagen (Advanced BioMatrix) [119]. The different treatments (indicated in the figures) were: proliferation and differentiation (24 h). Presence and absence of CuSO_4_ and/or Tetraethylenepentamine (TEPA); the concentrations used were 100 μM for proliferating cells and 30 μM for differentiating cells as previously described [36].

### Plasmid construction, virus production, and transduction of primary myoblasts

For shRNA viral production, Mission plasmids (Sigma) encoding for two different shRNA against *Mtf1* and a scramble (*scr*) are indicated in Table **S4.** CRISPR/Cas9 plasmid construction was performed by custom-designed of four sgRNAs to recognize the exon 1, exon 2, exon 4 and exon 8 of MTF1 mouse gene (Reference Sequence: NM_008636.4). Each gRNA consisted of 20 nucleotides complementary to the sequence that precedes a 5’-NGG protospacer-adjacent motif (PAM) located in the targeted exon. Specificity was validated by search through the entire genome to avoid off-target effects. Preparation of CRISPR/Cas9 lentiviral constructs was performed using the lentiCRISPRv2 oligo cloning protocol [120]. Briefly, sense and antisense oligos obtained from Integrated DNA technology (IDT), were set according to the designed gRNA and were annealed and phosphorylated to form double stranded oligos. Subsequently, they were cloned into the BsmBI–BsmBI sites downstream from the human U6 promoter of the lentiCRISPRv2 plasmid [120, 121] that was a kind gift from Dr. Feng Zhang (Addgene plasmid # 52961). The empty plasmid that expresses only Cas9 but no gRNA was included as null knock out control. The ligation reaction was used to transform competent Stbl3 bacteria, and positive cloning was confirmed by sequencing. Oligonucleotides used to form double stranded gRNAs are listed in **Table S5.**

To generate Lentiviral particles, 5 × 10^6^ HEK293T cells were plated in 10 cm dishes. The next day, transfection was performed using 15 μg of each shRNA and the packing vectors pLP1 (15 μg), pLP2 (6 μg), pSVGV (3 μg). CRISPR/Cas9 lentiviral particles were generated using a mixture of 12 μg of each construct, 9 μg of psPAX2 and 3 μg of pVSV-G packaging vectors. Plasmid mistures were diluted in 3 ml OptiMEM (Life Technologies) and supplemented with 60 μl Lipofectamine 2000 (Invitrogen). The complete mixture was incubated for 15 min before being added to cells. After an overnight incubation, the media was changed to 10 ml DMEM (Life Technologies) with 10 % FBS (Life Technologies). The viral supernatant was harvested after 48 h of incubation and filtered through 0.45 μm syringe filter (Millipore). To infect primary myoblasts, 5 ml of the filtered supernatant supplemented with 8 μg/ml polybrene (Sigma) were used to infect two million cells for overnight. Infected cells were then selected in DMEM/F12 (Life Technologies) containing 20% FBS, 0.75 ng/ml of Fibroblast Growth Factor and 1.5 μg/ml puromycin (Invitrogen).

### Antibodies

Primary antibodies (used at 1:1000) were obtained from Santa Cruz Biotechnologies: rabbit anti-MTF1 (sc-365090), rabbit anti-PI3K (sc-515646), mouse anti-β-actin (sc-81178), mouse anti-SERCA (sc-271669), mouse anti-RNA Polymerase II (sc-271669). From Abclonal: rabbit anti-caspase 3 (A2156), rabbit anti-MyoD; and a rabbit anti-GAPDH-HRP was from Sigma (G9295). Normal l Rabbit IgG was obtained from Cell signalling Technologies (2729). The anti-myosin heavy chain (MF20, deposited by D. A. Fischman), anti-myogenin antibody (F5D, deposited by W.E. Wright) was obtained as hybridoma supernatants from the Developmental Studies Hybridoma Bank (University of Iowa). The secondary antibodies used were: goat anti-rabbit and anti-mouse coupled to HRP (1:1000, Life Technologies).

### Primary myoblast immunofluorescence

Primary myoblasts for immunofluorescence were grown on glass bottom Cellview Advanced TC culture dishes (Grenier Bio One). Samples were obtained for proliferation and at 24, 48 and 72 h after induction of differentiation. Cells were fixed in 10% formalin, permeabilized with PBT buffer (0.5% Triton-X100 in PBS) and blocked in 5% horse serum in PBT. Cells were incubated with the rabbit anti-MTF1 antibody (1:100) in blocking solution overnight at 4 °C. The samples were then washed three times with PBT solution for 10 min at room temperature. Then, the cells were incubated with the goat anti-rabbit Alexa-488 secondary antibody (1:500) in blocking solution for 2 h at room temperature and 30 min with DAPI. Cells were counterstained with DAPI and imaged with a Leica TCS SP5 Confocal Laser Scanning Microscope (Leica) using a 40X water immersion objective.

### Immunohistochemistry

Proliferating and differentiating primary myoblasts at the desired time points were fixed overnight in 10% formalin-PBS at 4 °C. Samples were washed with PBS and permeabilized for 10 min in PBS containing 0.2% Triton X-100. Immunohistochemistry was performed using universal ABC kit and developed with Vectastain Elite ABC HRP kit (Vector Labs) following manufacturer’s instructions.

### Gene expression analyses

Three independent biological replicates of proliferating and differentiating (24 h) primary myoblasts were washed with ice cold PBS and RNA extracted using Trizol (Invitrogen). cDNA synthesis was performed using 1 μg of RNA, DNase I amplification grade (Invitrogen 18068-015) and Superscript III (Invitrogen 18080-400) according to manufacturer’s instructions. Changes in gene expression were analyzed by quantitative RT-PCR using Fast SYBR-Green master mix (Thermofisher Scientific) on the ABI StepOne Plus Sequence Detection System (Applied Biosystems) using the comparative Ct method [122] using *Ef1α* as control. The primers are listed in **Table S5**.

### Chromatin Immunoprecipitation (ChIP) Assays

Three independent biological replicates of proliferating and differentiating (24 h) primary myoblasts were crosslinked with 1% Formaldehyde and incubated for 10 min at room temperature on an orbital shaker. To inactivate the formaldehyde, 1 ml of 1 M glycine was added and cells were incubated for 5 min on an orbital shaker at room temperature. Cells were washed 3 times with 10 ml of ice cold PBS supplemented with Complete Protease

Inhibitor (Roche). Crosslinked myoblasts were resuspended in 1 ml of ice cold PBS supplemented with Complete Protease Inhibitor. The cell suspension was centrifuged for 5 min at 5,000 x *g* at 4 °C. The PBS was removed and the cell pellet was resuspended in 200 μl of ice cold SDS lysis buffer (50 mM Tris pH 8; 10 mM EDTA, 1% SDS) for 10 min. Proliferating myoblasts were sonicated three times for 5 min, 30” by 30” at mild intensity for myoblasts and five times for nascent myotubes using a Bioruptor UCD-200 (Diagenode). The samples were diluted to a final volume of 1 ml in ChIP buffer (16 mM Tris pH 8.1; 1.2 mM EDTA; 0.01% SDS; 1.1% Triton X100; 167 mM NaCl). ChIP was performed using a rabbit anti-MTF1 and a rabbit IgG antibodies. Samples were incubated for 2 h at 4 °C in a rotating platform and subsequently, 80 μl of Magna ChIP protein A+G Magnetic Beads (Millipore) were added to each sample and incubated overnight in a rotating platform at 4 °C. The samples were then placed in a magnetic rack and sequentially washed using 1 ml each of the wash buffer sequence A-D (Buffer A: 20 mM Tris pH 8.1, 2 mM EDTA, 0.1% SDS, 1% Triton-X100, 167 NaCl, Buffer B: 20 mM Tris pH 8.1, 2 mM EDTA, 0.1% SDS, 1% Triton-X100, 500 NaCl; Buffer C: 10 mM Tris pH 8.1, 1 mM EDTA, 1% NP40, 1% Sodium deoxicholate, 0.25 M LiCl_2_; Buffer D: 10 mM Tris pH 8.1, 1 mM EDTA). Immune complexes were eluted in 100 μl of buffer containing 0.1 M NaHCO_3_, 1% SDS, 1 μg/μl proteinase K. Samples were then reverse-crosslinked by adding 20 μl of 5 M NaCl and incubating overnight at 65 °C. The reverse crosslinked DNA was purified using the ChIP DNA clean concentrator, following the manufacturer’s instructions (Zymo Research). The DNA was stored at −80 °C until further analysis by semi-quantitative real time PCR or library preparation for ChIP-Seq.

### ChIP-seq

#### Library construction

Libraries of ChIP-enriched DNA were prepared from two biological replicates following the Illumina strategy. Samples were end-repaired, A-tailed, and adaptor-ligated using barcoded inline adaptors according to the manufacturer’s instructions (Illumina). DNA was purified over a Zymo Research PCR purification column between each enzymatic reaction. DNA was PCR amplified with KAPA HiFi polymerase using 16 cycles of PCR. Each library was size-selected for 200-300 bp fragments on a 1.5% agarose gel and the library concentrations were determined using a QuBit 3.0 Fluorometer (Thermo Scientific). Libraries were sequenced on an Illumina HiSeq2000 using single-end 50 bp sequencing at the University of Massachusetts Medical School deep sequencing core facility.

#### Data Analysis

Single-end fastq reads were split by barcode adapter sequences and adapter sequences were removed using the Fastx toolkit. Reads were mapped to the mm10 genome using bowtie, allowing up to three mismatches. Aligned reads were processed using HOMER [123]. UCSC genome browser tracks were generated using the “makeUCSCfile” command”. Mapped reads were aligned over all annotated mm10 TSSs using the “annotatePeaks” command, generating 20 bp bins and summing the reads within each window. After anchoring mapped reads over reference TSSs, aggregation plots were generated by averaging data obtained from two biological replicates. Peaks were called individually from replicate datasets using the “findPeaks” command and then overlapping peaks were identified using the “mergePeaks” command. For peak calling, a false discovery rate (FDR) of 0.001 was used as a threshold. Motifs were identified using the “findMotifs” command.

Analysis of data from GSE24852 [106] was performed similarly. Data was downloaded from GSE24852 and converted to fastq files using SRAtoolkit fastq-dump and mapped reads were converted to mm10. Aligned reads were processed in HOMER [123], as described above.

#### Gene Ontology (GO) term identification

GO term analysis was performed on metascape http://metascape.org [124].

### Sequential chromatin immunoprecipitation (Re-ChIP)

Primary myoblasts were lysed using the SimpleChIP^®^ Plus Sonication Chromatin IP Kit (Cell Signaling Technologies), following the manufacturer’s instructions. Briefly, after incubating the samples with MTF1 antibody and collecting immunoprecipitated material with magnetic beads, the samples were incubated with an equal volume of 10 mM dithiothreitol (DTT) for 30 min at 37 °C [125–127]. The supernatant was used for the second immunoprecipitation by adding a rabbit anti-MyoD antibody and incubating the samples similarly to the first immunoprecipitation. IgG substituted for the MTF1 and MyoD antibodies served as a negative control.

### Western blot analysis

Proliferating and differentiating primary myoblasts were washed with PBS and solubilized with RIPA buffer (10 mM PIPES, pH 7.4, 150 mM NaCl, 2 mM EDTA, 1% Triton X-100, 0.5% sodium deoxycholate, and 10% glycerol) containing Complete Protease Inhibitor. Protein was quantified by Bradford [128]. Samples (20 μg) were prepared for SDS-PAGE by boiling in Laemmli buffer. The resolved proteins were electrotransferred to PVDF membranes (Bio-Rad). The proteins of interest were detected with the specific polyclonal or monoclonal antibodies. Then the membranes were incubated for 2 h at room temperature with the species-appropriate peroxidase-conjugated antibodies (Invitrogen). Chemiluminescent detection was performed with ECL PLUS (GE Healthcare). Experiments were performed using samples from three independent biological experiments.

### Immunoprecipitation

Cells were washed three times with ice-cold PBS and resuspended in IP lysis buffer (50 mM Tris-HCl, pH 7.5, 150 mM NaCl, 1% Nonidet P-40, 0.5% sodium deoxycholate, and Complete Protease Inhibitor). Cell extracts were incubated with the anti-MTF1 primary antibody at 4 °C for 2 h, followed by an overnight incubation with PureProteome Protein A/G mix magnetic beads (Millipore). Samples were washed as indicated by the manufacturer, and immunoprecipitated proteins were eluted in freshly prepared IP-elution buffer (10% glycerol, 50mM Tris HCl, pH 6.8, and 1 M NaCl) at room temperature for 1 h [42]. Samples were analyzed by SDS-PAGE and Western blot.

### Subcellular fractionation of primary myoblasts and metal content analysis

Three independent biological replicates of proliferating and differentiating (24 h) primary myoblasts were fractionated using the Rapid Efficient And Practical (REAP) nuclear and cytoplasmic separation method [129]. Briefly, cells were washed with ice cold PBS, scraped and transferred to a 1.5 ml microcentrifuge tube. Samples were centrifuged for 10 seconds at 13,000 x *g* and the supernatant was discarded. The samples were resuspended in 500 μl of ice cold PBS containing 0.1% NP40 (Calbiochem) and 100 μl of the cell suspension were collected as the whole cell fraction. The remaining 400 μl were used to obtain nuclear and cytosolic fractions by disrupting the cells by pipetting using a 1 ml pipette tip. Cell suspension was centrifuged for another 10 sec and the supernatant was collected as the cytosolic fraction. The nuclear pellet was then washed twice in 1 ml of ice cold PBS containing 0.1% NP40 and once again centrifuged for additional 10 sec. The supernatant was removed and pellet was resuspended in 100 μl of PBS. Nuclear integrity was verified by light microscopy. All samples were sonicated at medium intensity for 5 min in 30 s on 30 s off cycles. Protein was quantified by Bradford method [128]. Purity of the fractions was evaluated by western blot.

The comparative analysis of copper concentrations from each sample was carried out using an AAS equipped with a graphite furnace (PerkinElmer, AAnalyst 800). A known mass of sample was acid digested in concentrated HNO_3_, using a single-stage digestion method [130, 131]. All measurements were performed in triplicate, resulting in a limit of detection for Cu of 15 ppb, calculated as 3σ. Analytical grade standards were used and diluted in 18 MΩ purified water. Copper content on each sample was normalized to the initial mass of protein.

### Expression and purification of recombinant MTF1

MTF1 wild type pET-GST/TEV/mMtf1[NM_008636.4] plasmid was purchased from Vector Builder. This plasmid was used as a template to introduce the mutations coding for the multiple Alanine substitutions in the putative carboxy-terminus Cu^+^-binding site (MBS) using the oligos indicated in **Table S5**. Mutations were introduced using the quick change mutagenesis kit following the manufacturer’s instructions (Agilent Technology). The pET vector places a GST tag at the amino-terminus, which was used for purification of the recombinant proteins. DNA sequences were confirmed by automated sequencing.

Plasmids coding for MTF1 wild type and the MBS mutant proteins were transformed into Stbl3 cells for propagation and transformed into BL21 DE3 bacteria for expression. Recombinant protein expression was performed according to an auto-inducing medium protocol [132]. Purification of GST-tagged wild type and mutated MTF1 recombinant proteins was carried out using Glutathione Agarose resin as described by the manufacturer (Pierce). Purified proteins were stored at −20 °C in Buffer containing 10% glycerol, 100 mM Tris, pH 8, and 150 mM NaCl. Protein concentrations were determined by Bradford assay [128]. Molar protein concentrations were estimated using MW 81,000 Da for both MTF1 proteins. In order to eliminate any bound metal, all purified proteins were treated with metal chelators as described previously [133–136]. Briefly, the proteins were incubated for 45 min at room temperature with 0.5 mM EDTA and 0.5 mM tetrathiomolybdate. Chelators were removed by buffer exchange using either 50 kDa cut-off Centricons (Millipore). The final purity of all protein preparations was at least 95%, as verified by SDS-PAGE followed by Coomassie Brilliant Blue staining and Western blot.

### Cu Loading to MTF1 and metal binding analyses

Cu^+^ loading was performed by incubating each apo-protein (10 μM) in the presence of 10 M excess of CuSO_4_, 25mM Hepes (pH 8.0), 150 mM NaCl and 10 mM ascorbate for 10 min at room temperature with gentle agitation, as previously described [136]. The unbound Cu was removed by washing in 50 kDa cut-off Centricons. Levels of Cu bound were verified by AAS, Varian. Briefly, before determinations, sample aliquots were mineralized with 35% HNO3 (trace metal grade) for 1 h at 80 °C, and digestions were concluded by making the reaction 3% H_2_O_2_. Metal bound to wild type and mutant MTF1 was measured in triplicate using a method similar to the subcellular fractionation of primary myoblasts (*vide supra*).

### Statistical analyses

In all cases, the data represent the average of three independent biological replicates ± SE. Two-tailed t tests were performed for statistical analyses using Graphpad Prism, Version 7

### Data availability

Genomic datasets have been deposited within GEO.

## Acknowledgements

We thank Ms. Courtney McCann, Dr. Sabriya Syed, Dr. Hanna Witwicka, and members of the Padilla-Benavides lab for insightful discussions regarding this manuscript. This work was supported by the Faculty Diversity Scholars Award from the University of Massachusetts Medical School to TP-B and by funding from the National Institute of Health (R01GM56244 to ANI and R01HD072122 to TGF) and from the National Science Foundation (DBI 0959476 to JGN). S.J.H. is a Special Fellow of the Leukemia and Lymphoma Society.

## Author Contributions

Conception and design: T.P.-B.; Acquisition of data: C.T.-M., S.J.H., D.C., S.J.V.G., Y.X., P.R.-G. J.G.N. AND T.P.-B.; All the authors were involved in the analysis and interpretation of data, as well as in drafting and revising the article.

