## Supplemental information for "MTF1, a classic metal sensing transcription factor, promotes myogenesis in response to copper"

Short title: MTF1 promotes skeletal muscle differentiation

### SUPPORTING INFORMATION

**SUPPORTING FIGURES**

**Figure S1**

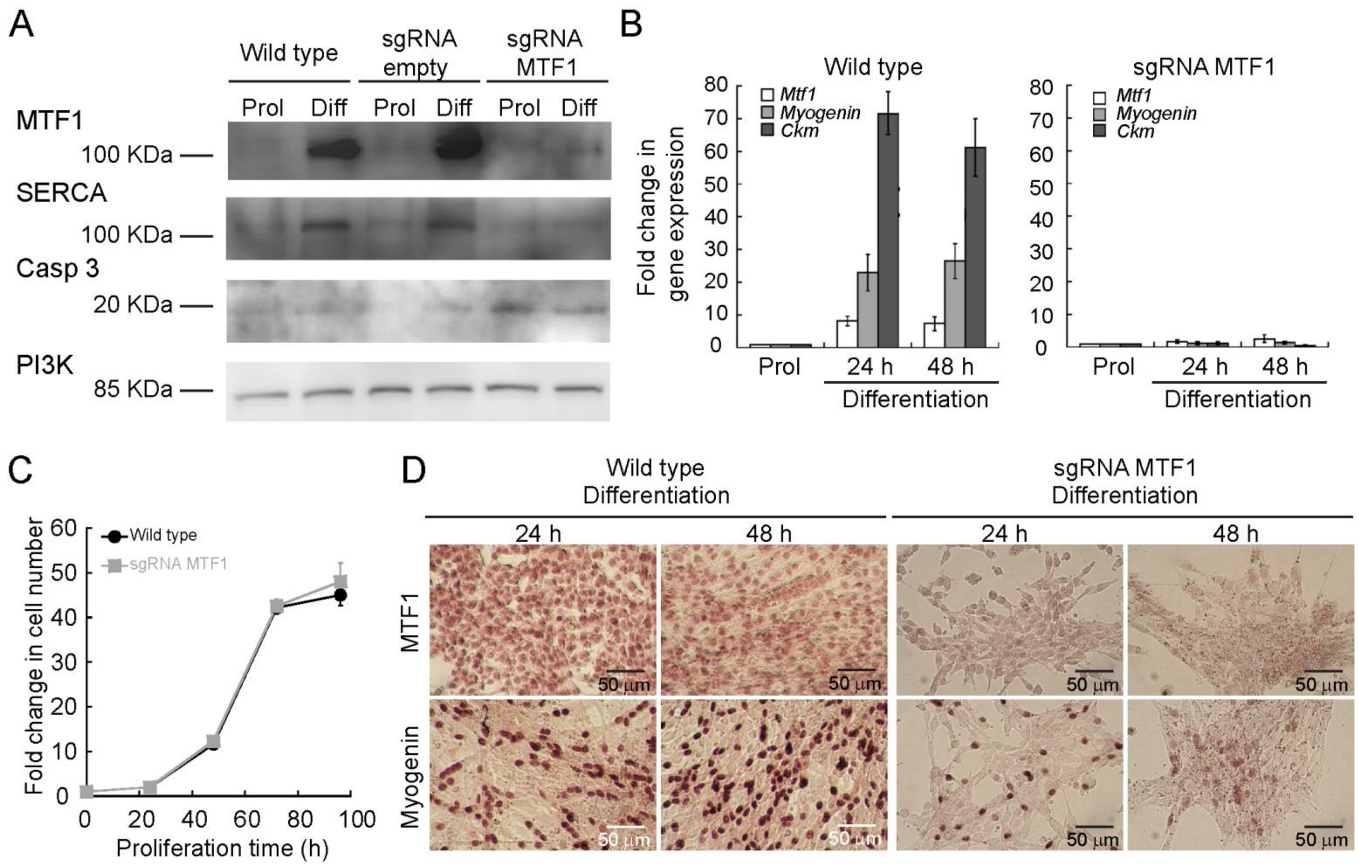


***Figure S1. Partial depletion of MTF1 using CRISPR/Cas9 inhibits myoblast differentiation, but does not affect proliferating cells.*** (A) Representative western blot of proliferating and differentiating wild type, empty sgRNA, and MTF1 partially depleted primary myoblasts. Levels of MTF1 and the differentiation markers myogenin and SERCA are decreased in myoblasts transduced with the sgRNA against MTF1, while the apoptosis marker Caspase 3 is increased in these cells. PI3K was used as loading control. (B) Steady-state mRNA levels of *Mtf1*, *Myogenin* and *muscle specific creatine kinase* (*Ckm*) in proliferating and differentiating primary myoblasts determined by qRT-PCR and normalized to *Ef1α*. Data represent the average of three independent biological replicates ± SE. (C) Proliferation curves comparing wild type and MTF1 partially depleted primary myoblasts. Data represent the average of three independent experiments ± SE. (D) Representative light micrographs of differentiating myoblasts were immunostained for MTF1 (upper panel), and myogenin (lower panel).

### Figure S2

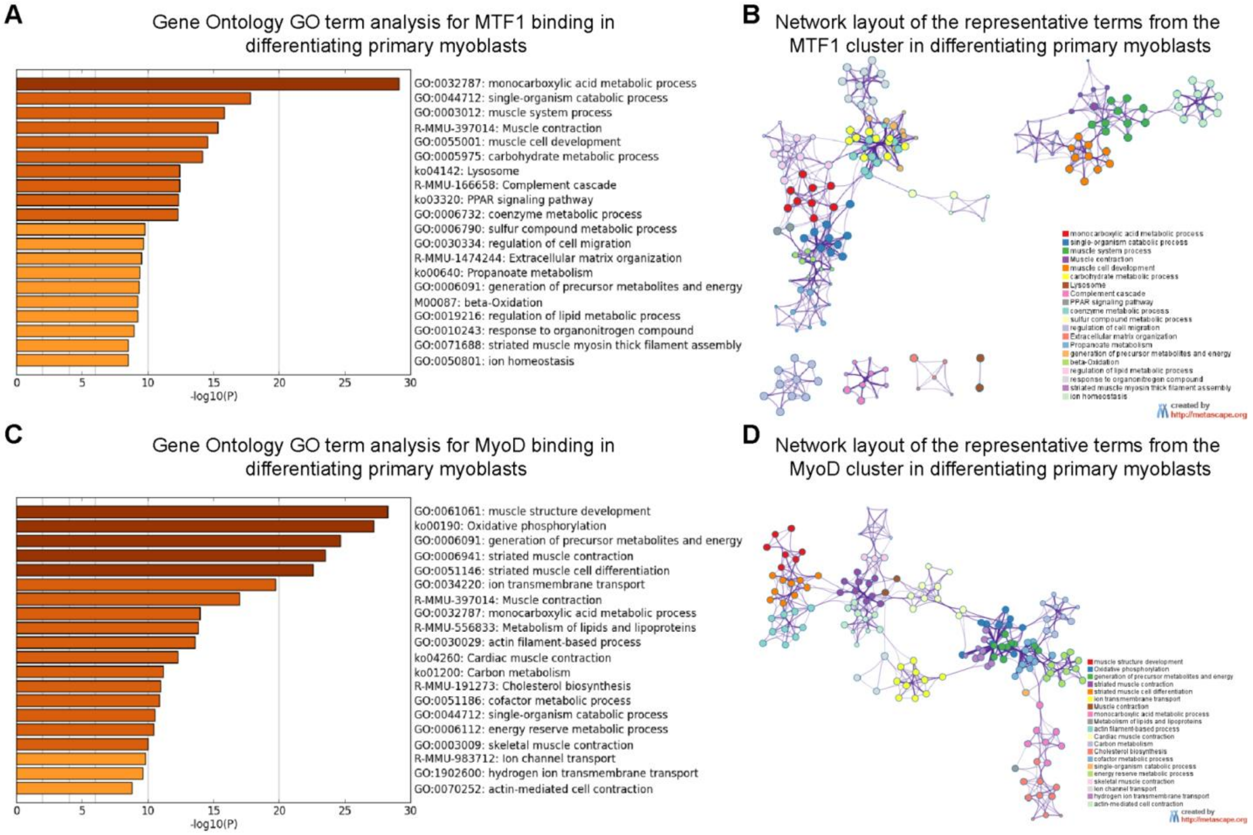


***Figure S2. GO analysis of MTF1/MyoD annotated peaks under differentiation conditions***. Heatmaps of enriched terms across MTF1-bound genes (A) or MyoD-bound genes (C) identified in differentiated primary myoblasts. Protein-protein interaction enrichment analysis for GO terms enriched in MTF1-bound genes (B) or MyoD-bound genes (D). The networks were generated using metascape [1], which utilizes BioGrid [2] and molecular complex detection (MCODE) algorithm [3] to identify network connections.

### Figure S3

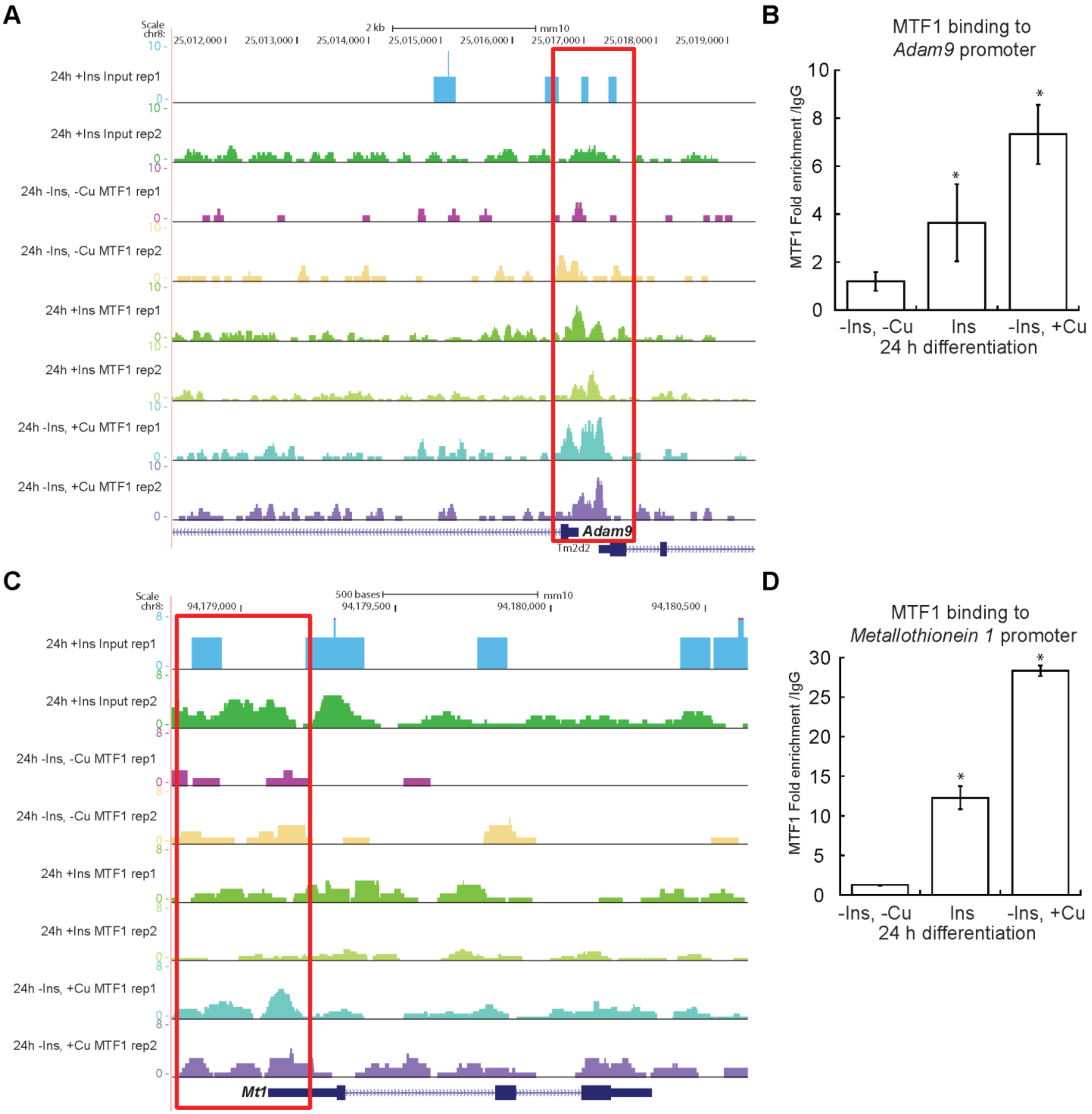


***Figure S3. MTF1 binds to additional myogenic genes promoters like Adam9 and to classic targets such as Metallothionein 1***. Genome browser tracks of replicate ChIP-seq and quantitation of ChIP-qPCR experiments examining endogenous MTF1 occupancy in differentiating myoblasts (24h) cultured under different concentrations of Cu. Additional example of MTF1 binding to a target myogenic gene *Adam9* (A, B). *Metallothionein 1* (*Mt1*) is the classic metal homeostasis gene target of MTF1 and was used as positive control (C, D). ChIP-qPCR data represent the average of three independent experiments ± SE.

**Figure S4**

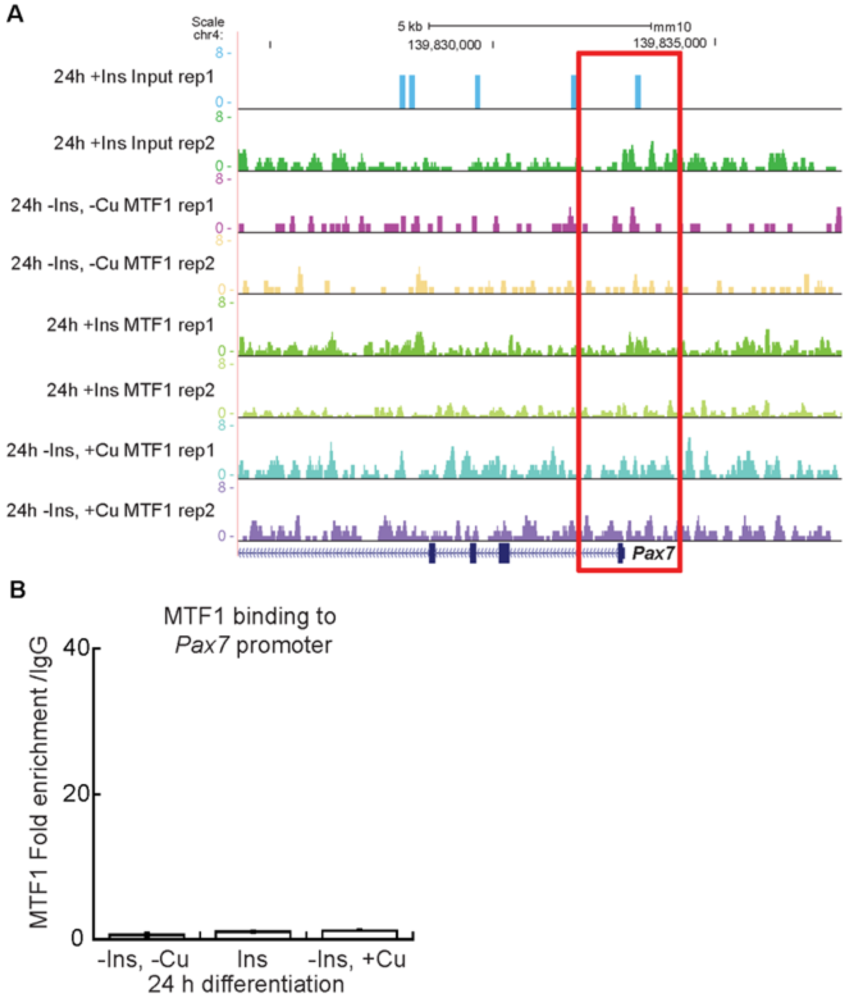


***Figure S4. MTF1 does not bind to the Pax7 promoter***. Genome browser tracks of replicate ChIP-seq (A) and quantitation of ChIP-qPCR (B) experiments examining endogenous MTF1 occupancy to the *Pax7* promoter in differentiating myoblasts (24h) cultured under different concentrations of Cu. ChIP-qPCR data represent the average of three independent experiments ± SE.

**Figure S5**

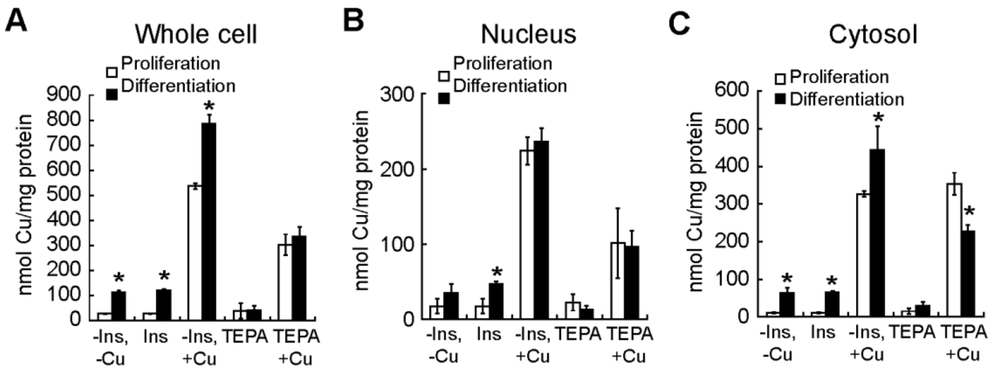


***Figure S5. Cellular distribution of Copper in differentiating myoblasts.*** (A) Whole cell Cu content of proliferating and differentiating primary myoblasts determined by AAS. Nuclear (B) and cytosolic (C) Cu content of proliferating and differentiating primary myoblasts cultured under different Cu conditions. Metal determination was performed by AAS. Data represents the mean Cu concentration of three independent biological replicates ± SE. *P < 0.01.

**SUPPLEMENTAL TABLES**

**Supplemental table 4. shRNA Mission plasmids (Sigma) used in this study**

| Oligo name | Sequence 5’→3’ | Catalog No. |
|---|---|---|
| *Mtf1* shRNA-1 | CCGGGCTCACCTTTCAGACGTATTTCTCGAGAAATAC GTCTGAAAGGTGAGCTTTTTG | TRCN0000312828 |
| *Mtf1* shRNA-2 | CCGGGCCAACTCTGTCCTAACTAATCTCGAGATTAGT TAGGACAGAGTTGGCTTTTTG | TRCN0000312778 |

**Supplemental table 5. Oligos used in this study**

| Oligo name | Sense 5’→3’ | Antisense 5’→3’ | Primer use |
|---|---|---|---|
| MTF1 gRNA1 | CACCGAAGAAGATGATATTGTCGTC | AAACGACGACAATATCATCTTCTTC | gRNAs for CRISPR Cas9 |
| MTF1 gRNA2 | CACCGGTACACCTTCGTCTGTAATC | AAACGATTACAGACGAAGGTGTACC | gRNAs for CRISPR Cas9 |
| MTF1 gRNA3 | CACCGACCACCTTAAAACTCACGTC | AAACGACGTGAGTTTTAAGGTGGTC | gRNAs for CRISPR Cas9 |
| MTF1 gRNA4 | CACCGGCCTTGATTGACAGTTTTAA | AAACTTAAAACTGTCAATCAAGGCC | gRNAs for CRISPR Cas9 |
| *Myogenin* promoter | ACGCCAACTGCTGGGTGCCA | GAATCACATGTAATCCACTGGA | ChIP-qPCR |
| *Adam9* promoter | CCGCCCAGCACCATCTTC | CTACTGGAGGGCGGAGCG | ChIP-qPCR |
| *Mt1* promoter | CTCCGCCCGAAAAGTGCGCTC | GAAGCTGGAGCTACGGAGTAA | ChIP-qPCR |
| *Pax7* promoter | GTGGCGACAAGGAAGTTCAAACAAAC | AAAGAAAGCCACTCCGCAACCTCTG | ChIP-qPCR |
| q*Myogenin* | CAAGTGTGCACATCTGTTCTAGTCTCT | GTATCATCAGCACAGGAGACCTTGGT | Gene expression qPCR |
| q*ckm* | GCCGGGGATGAGGAGTCCTAC | GCAGTGCGGAGGCAGAGTGTA | Gene expression qPCR |
| q*Mtf1* | AGATGATATTGTCGTCTGGACTGTG | GAAGCGGAAGTGACGCTAGGGACAG | Gene expression qPCR |
| q*Ef1α* | AGCTTCTCTGACTACCCTCCACTT | GACCGTTCTTCCACCACTGATT | Gene expression qPCR |
| MBS MTF1 | CCTTCGCAGAGTCCCGGGCCGCGGCCT GAGCCTGAGCGGCCTCCTCTTGTTTGA | TCAAACAAGAGGAGGCCGCTCAGGCTC AGGCCGCGGCCCGGGACTCTGCGAAGG | Site directed mutagenesis |

**SUPPLEMENTAL RESOURCES**

The following major datasets were generated

| Author (s) | Year | Dataset title |
|---|---|---|
| C. Tavera-Montañez, S.J. Hainer, D. Cangussu, S.J.V. Gordon, Y. Xiao, P. Reyes-Gutierrez, A. N. Imbalzano, J.G. Navea, T.G. Fazzio and T. Padilla-Benavides | 2019 | MTF1, a classic metal sensing transcription factor, promotes myogenesis in response to copper. |

The following previously published datasets were used

| Author (s) | Year | Dataset title | Dataset URL | Database, license, and accessibility information |
|---|---|---|---|---|
| Soleimani VD, Rudnicki MA | 2012 | ChIP-Seq of Myf5, MyoD, Snai1, HDAC1, HDAC2, E47 and empty vector controls in mouse skeletal myoblasts or myotubes | https://www.ncbi.nlm.nih.gov/geo/query/acc.cgi?acc=GSE24852 | Publicly available at the NCBI Gene Expression Omnibus (accession no: GSE24852) |
| Soleimani VD, Rudnicki MA | 2012 | MyoD-2DM | https://www.ncbi.nlm.nih.gov/geo/query/acc.cgi?acc=GSM611279 | Publicly available at the NCBI Gene Expression Omnibus (accession no: GSM611279) |
